# Genome fractionation and loss of heterozygosity in hybrids and polyploids: mechanisms, consequences for selection, and link to gene function

**DOI:** 10.1101/2020.07.30.229369

**Authors:** Karel Janko, Oldřich Bartoš, Jan Kočí, Jan Roslein, Edita Janková Drdová, Jan Kotusz, Jan Eisner, Eva Štefková-Kašparová

## Abstract

Hybridization and genome duplication have played crucial roles in the evolution of many animal and plant taxa. During their evolution, the subgenomes of parental species undergo considerable changes in hybrids and polyploids, which often selectively eliminate segments of one subgenome. However, the mechanisms underlying these changes are not well understood, particularly when the hybridization is linked with asexual reproduction that may enforce specific evolutionary pathways.

We studied the genome evolution in asexual diploid and polyploid hybrids between fish from the genus *Cobitis*. Comparing exome sequencing with published cytogenetic and RNAseq data revealed that clonal genomes remain static on chromosome-scale levels but undergo considerable small-scale restructurations owing to two major processes; hemizygous deletions and gene conversions. Interestingly, polyploids were much more tolerant to accumulating deletions than diploid asexuals where gene conversions prevailed. The genomic restructurations accumulated preferentially in genes characterized by high transcription levels, relatively strong purifying selection and some specific functions such as interacting with intracellular membranes. The likelihood of an ortholog’s retention or loss correlated with its parental-species ancestry, GC content, and expression. Furthermore, all hybrids showed a strong bias towards the retention of one parental subgenome. Contrary to expectations, however, the preferentially retained subgenome was not transcriptionally dominant as all hybrids were phenotypically more similar to the other parent.

The present study demonstrated that the fate of subgenomes in asexual hybrids and polyploids depends on the complex interplay of selection and several molecular mechanisms whose impact depends on ploidy, sequence composition, gene expression as well as parental ancestry.

## Introduction

The genome of a typical Metazoan possesses two alleles of each gene brought together by merging reduced gametes of two individuals belonging to the same species. However, these rules have often been alleviated as traces of whole genome duplications (WGD) and introgressive hybridizations have been documented in many taxa, vertebrates and humans included (Dehal and Boore 2005; Gittelman et al. 2016). Hybridization and polyploidization may cause serious problems, e.g., in transcription regulation leading to deregulation of transposable elements (Dion-Côté et al. 2014; Lien et al. 2016), but may also lead to creation of novel traits and acquisition of gene functions via sub-/neofunctionalization of duplicated genes (Yoo et al. 2014; Fridman 2015), potentially facilitating specialization to new niches (Madlung 2013).

Realizing their evolutionary significance and huge practical value to mankind (Mason and Batley 2015), research focused on hybridization and polyploidy intensified and revealed some prominent patterns. For instance, hybrid phenotypes may range from intermediate forms to transgressive expression of novel traits (Bell and Travis 2005; Yoo et al. 2014; Bartoš et al. 2019); however, often, one parental subgenome is more expressed than the other one, known as ***expression dominance*** (Yoo et al. 2014; Alexander-Webber et al. 2016; Cheng et al. 2018). Hybrid and polyploid genomes evolve dynamically and often lose orthologous genes from one or the other parental species in processes referred to as ***loss of heterozygosity*** (LOH), ***genome fractionation***, or ***rediploidization*** in polyploids (Yoo et al. 2014; Lien et al. 2016; Du et al. 2020). These processes are often considerably asymmetrical (Alexander-Webber et al. 2016), and it has been proposed that loss of alleles from the less expressed parental subgenome may cause less severe effects and may, therefore, be preferred by selection (Yoo et al. 2014). However, the situation is likely more complex as orthologs may also be lost or retained for proper dosage of molecular interactors and relative copy number of their gene products, i.e., selection for stoichiometry (Birchler and Veitia 2012). Hybrid populations may also selectively filter orthologous genes according to their adaptive value in a given environment (Gittelman et al. 2016; Lancaster et al. 2019; Smukowski Heil et al. 2019). Thus, despite the application of modern technologies, the question why some genes tend to be retained in heterozygous or duplicated states, whereas others are subjected to fractionation still represents a major evolutionary puzzle. It remains particularly unclear whether the aforementioned patterns are driven by case-specific mechanisms or whether independent lineages follow similar evolutionary trajectories (Soltis et al. 2010; Deans et al. 2015).

Such a gap in current knowledge partly results from taxonomic bias in knowledge, particularly toward plant species, as the incidence of hybridization and polyploidy have traditionally been underrated by zoologists. Moreover, direct tests for determining adaptive values of genomic rearrangements could be performed under laboratory conditions only, thereby focusing on rapidly reproducing organisms (Smukowski Heil et al. 2017; Lancaster et al. 2019) as events like LOH are rather rare (Dukić et al. 2019). For practical reasons, most available data are derived from natural hybrids and polyploids, making it difficult to discern the patterns that are direct consequences of genome merging and those that evolved subsequently. In addition, many polyploids are of hybrid origin, making it challenging to discern the effects that are inherent to polyploidy and those to hybridization itself. Finally, in, it is unclear how many investigated hybrid and/or allopolyploid taxa establish themselves in natural environments since any new form is rare at the time of its emergence and is, therefore, threatened by frequency-dependent mating disadvantage and backcrossing with dominating ancestral populations, i.e., the ***minority cytotype exclusion principle*** (Husband 2000).

It has thus been hypothesized that new strains may alleviate initial caveats by taking advantage of ***asexual reproduction***, since production of unreduced gametes offers immediate reproductive isolation and clonal multiplication of novel genotypes unencumbered by maintaining two sexes (Cunha et al. 2008; Choleva and Janko 2013; Hojsgaard and Hörandl 2015; Janko et al. 2018; Dubey et al. 2019). The perception of asexual organisms indeed changed among to current appreciation that they occur in all major eukaryotic clades (Schön et al. 2009) and form dominant components in some ecosystems (Kearney 2005; Hojsgaard and Hörandl 2015). The emergence of asexual reproduction is tightly linked to hybridization and polyploidy, reviewed in (Choleva and Janko 2013), and may represent an inherent stage of the speciation process, representing a special type of Bateson–Dobzhansky–Muller model (Janko et al. 2018). This paradigm shift coincides with increasing interest in the role of recombination modification in evolution (Thompson and Jiggins 2014; Ortiz-Barrientos et al. 2016), suggesting that understanding the evolutionary processes in asexual organisms may provide important insights into the mechanisms of genome evolution in general.

Unfortunately, there is no consensus on genomic consequences of asexuality. Clonal inheritance has been originally assumed to ensure stasis of genome with gradual accumulation of deleterious mutations (Muller 1964; Keightley and Otto 2006) and heterozygosity levels (Birky 1996; Mark Welch and Meselson 2000; Balloux et al. 2003) or modified dynamics of transposable elements (Hickey 1982). This view is currently challenged by indications of horizontal gene transfers in some asexual lineages (Gladyshev et al. 2008; Danchin et al. 2010) as well as by accumulating evidence that genomes may acquire aneuploidy or structural changes extremely quickly once the sex is lost, owing to relaxed constraints on the pairing of homologous chromosomes (Triantaphyllou 1981; Sunnucks et al. 1996; Normark 1999; Spence and Blackman 2000; Tucker et al. 2013). Heterozygosity may degrade quickly by hemizygous deletions and particularly by gene conversions (Tucker et al. 2013), which may lead to increased GC content in asexual genomes (Bast et al. 2018). Notably, the dynamics of asexual genome may also be determined by its mode of origin as nonhybrid asexuals appear to have lost most of their heterozygosity as a consequence of pervasive gene conversions, whereas hybrid asexuals generally express high levels of heterozygosity, probably indicating efficient clonal transmission of the parental genomes (Jaron et al. 2018).

With such varying patterns, it is difficult to discern the mechanisms that are taxon-specific and those that are related to asexual reproduction *per se*. A major complication is that the so-called asexual organisms form a very heterogeneous group by employing a wide spectrum of gametogenetic mechanisms, ranging from ameiotic processes (apomixis) to those involving more or less distorted meiotic divisions (automixis) (Stenberg and Saura 2009; Stenberg and Saura 2013). Some types of automixis involve intragenomic recombinations or exclusions of large genomic parts, thereby decreasing heterozygosity among the progeny (Bi and Bogart 2006), whereas other types are theoretically equivalent to apomixis. For example, many hybrid asexuals employ ***premeiotic endoduplication***, a mechanism wherein normal meiosis is preceded by WGD in oogonial cells. As a consequence, segregation and recombination presumably occur on bivalents between sister copies of the chromosomes rather than between the orthologs, resulting in clonal progeny (Lutes et al. 2010; Arai and Fujimoto 2013) (Fig. 1a).

**Fig. 1:**
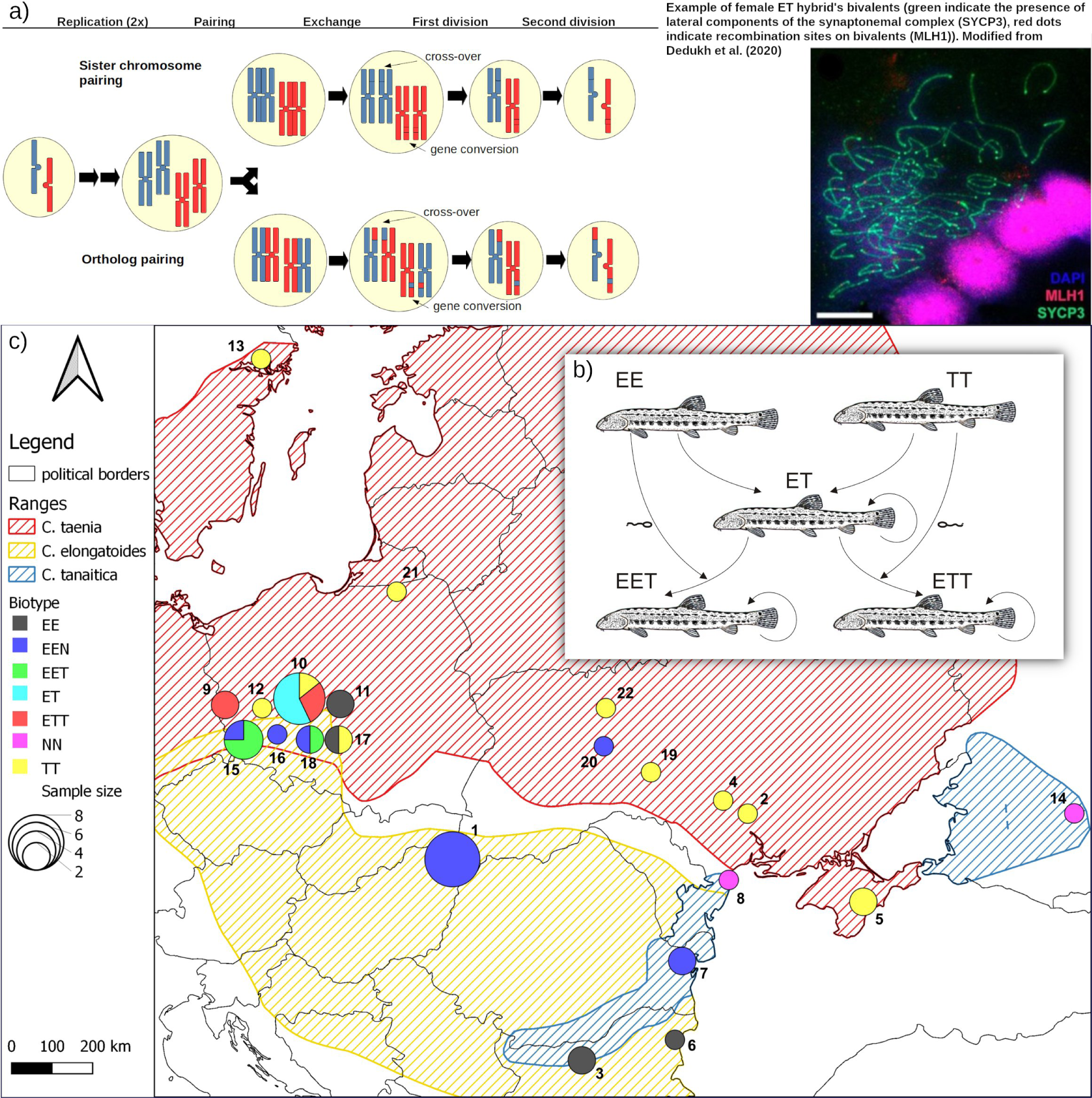
The *Cobitis taenia* hybrid complex. a) example of gametogenetic pathways involving endoreplication and followed by meiotic pairing of either sister chromosomes (upper pathway) or orthologous chromosomes (lower pathway). Note that the latter case may cause loss of heterozygosity among progeny via crossovers or gene conversions. Insert on the right demonstrates empirical evidence for the presence of proper bivalents in hybrid’s oocytes as from (Dedukh et al. 2020). b) Reproduction scheme of *Cobitis*; letters correspond to haploid genomes: E = *C. elongatoides*, T = *C. taenia*. c) Map of species distribution and samples’ origin; sites’ numeric code corresponds to Table S1.

In this study, we analyzed the causes and consequences of allelic recombination, conversion, and LOH in a clonally reproducing vertebrate of hybrid origin, *Cobitis* (Actinopterygii). We focused on the so-called *Cobitis taenia* hybrid complex, which arose by hybridization of the species *C. elongatoides* (we have denoted its haploid genome as “E”) with either of its two distant relatives, *C. taenia* (denoted as “T”) or *C. tanaitica* (denoted as “N”) (Choleva et al. 2012). Phylogenomic analysis (Janko et al. 2018) revealed that *C. taenia* diverged from *C. tanaitica* relatively recently, ca. 1 Mya (million years ago) but the initial E–(TN) divergence occurred ca. 9 Mya and was initially followed by intensive gene exchange. However, with ongoing divergence, these species lost the capacity to produce sexual hybrids as crossings of the current species led to sterile hybrid males but clonally reproducing, fertile hybrid females. Hybrid females form unreduced gametes by premeiotic endoduplication but are gynogenetic, i.e., they require sperm from a sexual species to trigger development of their gametes (Janko et al. 2007; Choleva et al. 2012; Dedukh et al. 2020). Usually, the sperm genome is degraded after fertilization, but a certain proportion of oocytes fuse with sperm cells, and consequently, diploid ET or EN females produce a certain portion of triploid progeny that may have EET, ETT, EEN, or ENN genomic constitution, depending on the sperm donor. Triploids also reproduce clonally (Fig. 1a, b).

Hybridization between the parental species is reciprocal, but *C. taenia* is the maternal ancestor of most *elongatoides*–*taenia* hybrids, whereas *C. elongatoides* is the maternal ancestor of *elongatoides*–*tanaitica* hybrids (Janko et al. 2003). Because parental species lack the obvious prezygotic reproductive barriers and glacial–interglacial cycles repeatedly brought their ranges into contact (Fig. 1c), new clones arose dynamically throughout much of the Pleistocene epoch and subsequently colonized Europe (Janko et al. 2005; Janko et al. 2012). Current asexual populations therefore consist of recent *elongatoides-taenia* clones with postglacial origin in the Central European hybrid zone, as well as of ancient *elongatoides*–*tanaitica* hybrids (the so-called hybrid clade I consisting of EN and EEN biotypes), which originated in the Balkan hybrid zone ca. 300 kya (kilo years ago) (Janko et al. 2005).

*Cobitis* hybrids, thus, offer excellent opportunity to investigate how genomes of natural asexual organisms evolve through time and across different ploidy levels. Majtánová et al. (2016) demonstrated by karyotypic analysis that clonal hybrids maintain remarkable integrity of the parental chromosomes without traces of large-scale recombinations and restructurations, despite more than 300 ky of evolution since the initial hybridization event. This study investigated the dynamics of diploid and polyploid clonal genomes on the finer scale of individual genes. To achieve this aim, we performed exome sequencing of the sexual parental species and their clonal hybrids and subsequently compared the data to recently published gene expression profiles (Bartoš et al. 2019). This allowed us to identify mechanisms underlying fractionation and LOH in clonal genomes and to test how they relate to expression and function of the affected genes.

## 3. Results

### 3.1. Polymorphism detection and identification of species-specific variants

Exome-capture data were acquired from 46 specimens, including three parental species and their asexual hybrids, which were selected to cover all major phylogroups, ploidy types and hybrid genome compositions (see Fig. 1c and Table S1 for details) and we also included whole genome sequencing of one ET hybrid for control. Reads were mapped against published *C. taenia* reference transcriptome containing 20,600 contigs (Janko et al. 2018) and for simplicity, we restricted our analysis to 189,927 quality filtered bi-allelic SNPs (i.e. single nucleotide polymorphisms occurring in no more than two states across the entire dataset). The greatest genetic divergence was between *C. elongatoides* and the remaining two sister species, *C. taenia* and *C. tanaitica*, while hybrids appeared intermediate (multidimensional scaling (MDS), Fig. 2a). Examination of pairwise genetic distances between hybrid individuals revealed 11 clusters of individuals that were nonrandomly similar to each other, therefore likely representing clonal lineages. We thus refer to them as independent Multilocus Lineages (MLL) *sensu* (Arnaud-Haond et al. 2007), see Fig. 2a and Table S2.

**Fig. 2:**
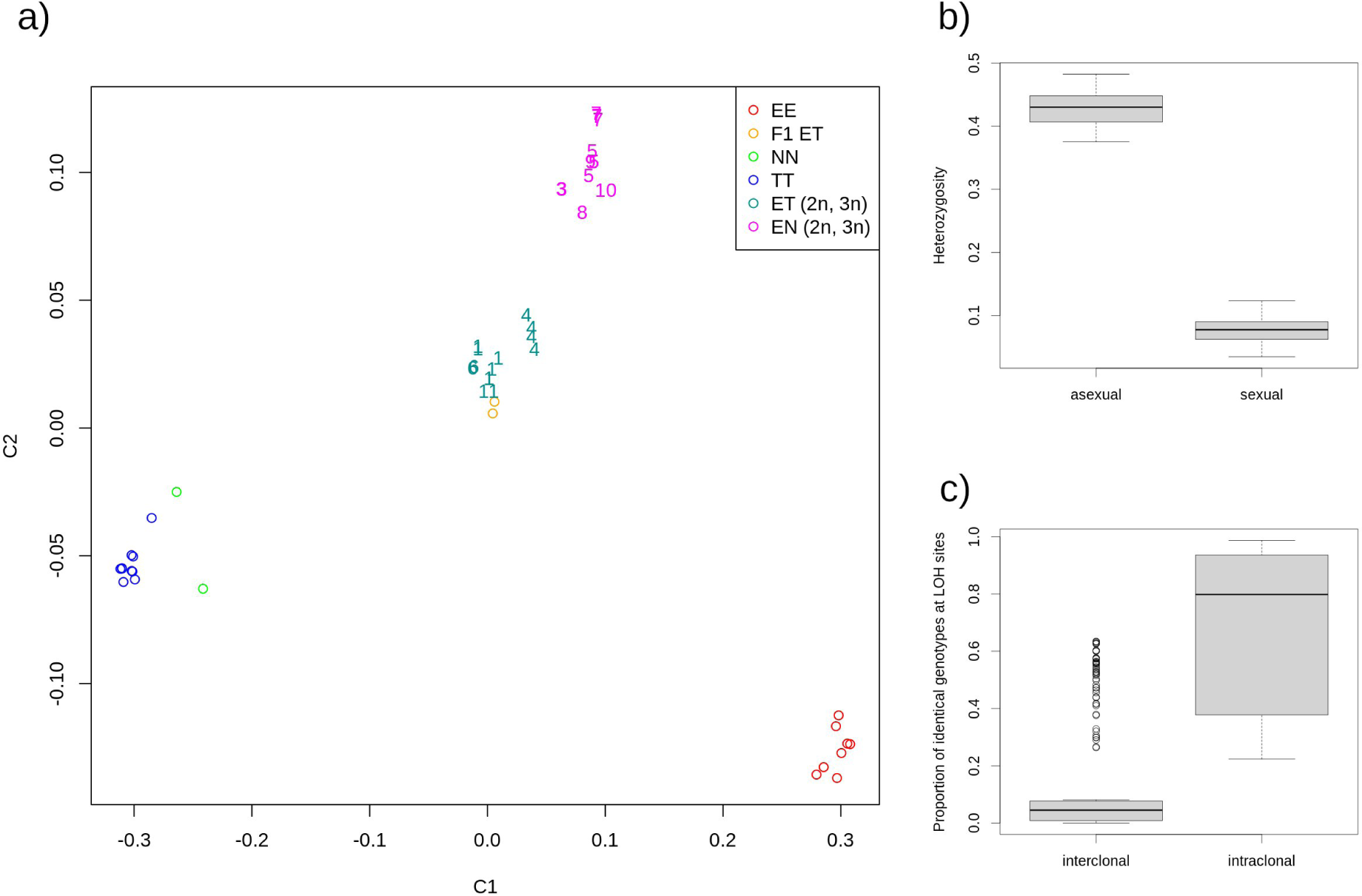
a) Multidimensional scaling (MDS ; SVD algorithm) of samples based on filtered recalibrated SNPs; clustering visualization of first two coordinates (Plink v1.9b) validates samples’ genetic origin; letters in the legend correspond to haplotype genomes as follows: *E* = *C. elongatoides*, *N* = *C. tanaitica*, *T* = *C. taenia*. Hybrid individuals are denoted numerals, which indicate the ID of determined clonal lineages, as in Table S2 (note that clone-mates tend to be clustered in the MDS plot). b) Heterozygosity of asexual and sexual samples across all filtered variant sites; missing sites are not included. c) Boxplots indicate the proportions of shared genotypes at LOH positions among all pairs of individuals belonging to the same (right) and different (left) clonal lineages. Note that we used the set of E-TN diagnostic sites to maintain compatibility of *elongatoides*-*taenia* and *elongatoides*-*tanaitica* hybrids.

To characterize hybrid’s SNP variation relative to their parental species, we followed (Ament-Velásquez et al. 2016) and divided all hybrids’ SNPs into 10 categories (Table S2), of which five were particularly important for this paper. First, we detected 16,372 unique positions with variants occurring only among asexuals but not among sexuals (SNP categories “***private-asexual*** 1a and 1b”). These SNPs presumably represent mutations acquired after clonal origin (Ament-Velásquez et al. 2016; Kočí et al. 2020). Notably, the proportions of such SNPs in genome of each hybrid were tightly correlated with its mtDNA distance from the nearest sexual counterpart (Pearson’s r=0.971, df=23, p-value=8.66e-16), so that the ancient *elongatoides*-*tanaitica* hybrids had the highest amount of private asexual SNPs, while experimental F1 hybrids had the least amount of such SNPs, probably representing only rare sequencing errors. Second, we focused on SNP variants that were diagnostic between pairs of parental species, so that their hybrids should possess one or both parental variants, thereby allowing detection of LOH events. Throughout the entire dataset we therefore identified sites diagnosing *C. elongatoides* from *C. taenia* (referred to as E-T diagnostic sites; total of 37,988), *C. elongatoides* from *C. tanaitica* (E-N diagnostic sites; 30,281) and we also found SNPs differentiating *C. elongatoides* from the joint dataset of *C. taenia* and *C. tanaitica* (E-TN diagnostic sites; 27,311). According to the way how hybrids’ SNPs were shared with these parental variants, we categorized them as “shared SNPs” of type sh3a (heterozygous for both parental variants), sh3b11 (homozygous for one parent’s allele) and sh3b12 (homozygous for the other parent’s allele) in Table S2.

### 3.2. Clonal lineages accumulate Loss of heterozygosity events in their evolution

Hybrids were considerably more heterozygous than parental species (Fig. 2b; Wilcoxon rank sum test: W = 520, p-value < 1e-9) with no less than 98.5% of private asexual SNPs and the vast majority of diagnostic sites occurring in heterozygous states. Nevertheless, LOH was observed in some portion of diagnostic SNPs of every hybrid (categories sh3b11 and sh3b12 in Table S2). We verified the quality of base-calling and LOH detection by two approaches.

We first compared SNP calling from exome-capture technology and whole-genome sequencing of the same ET hybrid (csc067) and found differences in only ∼0.17 % of E-T diagnostic positions. We also compared two F1 hybrids against their parents and found homozygous states only in ∼0.16 % of positions, where both parental individuals differed from each other. As these variants were suspiciously present in most of the other specimens, suggesting potential sequencing or demultiplexing errors rather than real variants, we excluded them form subsequent analyses. Overall, this indicates ***high reliability of LOH detection based on exome capture*.**

Two patterns were noted in the distribution of LOH SNPs. First, individual LOH sites were significantly more likely to be shared by individuals belonging to the same clonal lineage (MLL) than by individuals from different clones (Wilcoxon rank sum test, W = 4540, p-value < 1e-9; Fig. 2c). Second, the proportion of LOH sites in each individual significantly correlated with its proportion of private asexual mutations (Pearson’s r = 0.955, 95% c.i. = 0.902-0.980, p-value = 3.286e-14; Fig. 3a, Table S2). Hence, although we may not rule out existence of somatic mutations (López and Palumbi 2019), our data indicate that ***erosion of heterozygosity is heritable within clonal lineages, accumulates over clone’s evolutionary history, and therefore affects the germline*.**

**Fig. 3:**
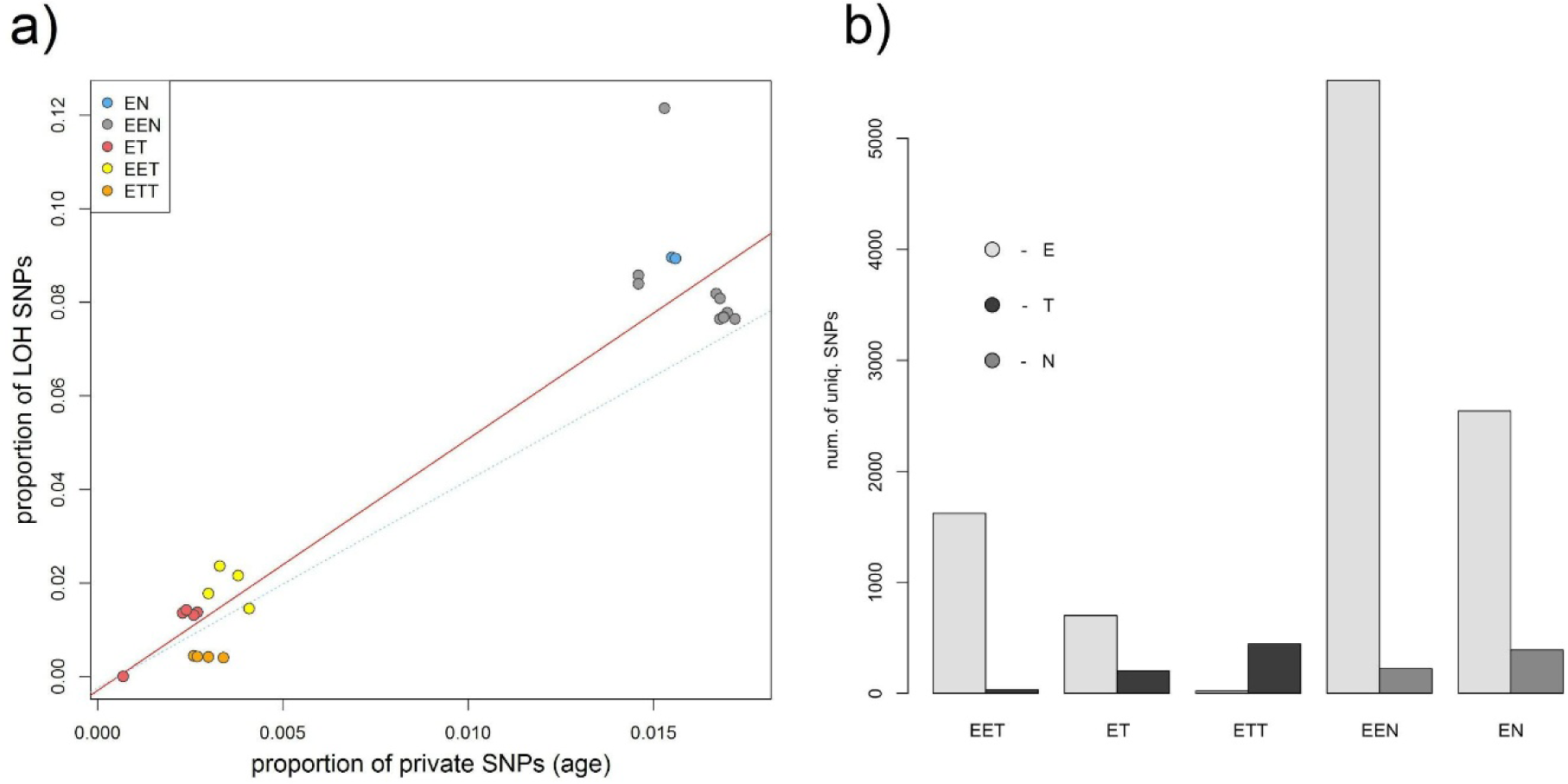
**a)** Correlation between proportion of private hybrid SNPs as the proxy for asexual’s age (x-axis), and proportion of LOH loci (y-axis) in each individual. For plotting points and their linear regression (solid line) we used E-T diagnostic sites for *elongatoides*-*taenia* hybrids and E-N diagnostic sites for *elongatoides*-*tanaitica* hybrids. Dashed line represents the linear regression fit for data calculated on E-TN diagnostic sites used for all individuals (points not shown). Letters correspond to haplotype genomes as follows: E = *C. elongatoides*, N = *C. tanaitica*, T = *C. taenia*. **b)** Barplot showing proportions of LOH events according to their genomic origin. Height of each bar represents absolute number of unique SNPs that appeared as LOH in a given biotype (note that barplots are not corrected for sample size of respective biotypes).

We also performed the same analysis on E-TN diagnostic sites, in order to minimize potential effect of ancestral polymorphism when same allele might have been inherited from both parents at time of clonal origin but was subsequently lost in one parental species. Since *C. taenia* – *C. tanaitica* divergence predates origin of the oldest *Cobitis* clones by hundreds of thousands of years (Janko et al. 2018), it may be posited that the most of such E-TN sites became diagnostic long before the origin of studied clones. Yet, proportions of LOH at E-TN diagnostic sites also correlated significantly with the private asexual SNPs (r = 0.948, 95% c.i. = 0.885-0.977, p-value = 2.145e-13) albeit with slightly less steep slope than in ET- and E-N sites, respectively (Fig. 3a). This suggests that some false-positives might have affected our dataset, but altogether the ***retention of ancestral polymorphism is an unlikely explanation of most observed LOH events***.

### 3.3. Heterozygosity deteriorates by gene conversions and hemizygous deletions in asexuals

We next investigated topological context of LOH sites to test if they might have been generated by point mutations. Since we used cDNA reference that excludes introns, we could not simply analyze the physical distance between SNPs in studies loci. Instead, we tested if LOH sites within individual genes tend to occur in contiguous stretches. We thus compared observed distribution of LOH sites to permuted datasets with LOH sites randomly distributed across all genes (see the Methods for details). We observed that numbers of LOH occurring in contiguous stretches significantly exceeded simulated values, suggesting that LOH sites tend to occur in clusters. This suggests that most ***LOH events are created by processes like gene conversions and deletions that affecticontiguous stretches of DNA***.

To distinguish between both candidate processes, we analyzed the sequencing coverage following (Tucker et al. 2013), who showed that conversions conserve the amount of allelic copies, while allelic deletions would result in coverage drop. To do so, we first provided a ***formal proof that exome capture data are suitable for coverage comparisons among individuals and across loci*** (Bragg et al. 2016) (see Appendix S1).We than used normalized per-SNP coverages to calculate relative values of coverage for each hybrids’ LOH by comparing it to the coverages of the same site in parental species (see Methods).

Following (Tucker et al. 2013) we predicted that conversions result in relative coverage ∼1, while allelic deletions would result in coverage drop to values ∼0.5 in diploids, or ∼0.66 (single deletion) and ∼0.33 (double deletion) in triploids. To investigate roles of both processes in LOH creation, we constructed for each hybrid biotype the histograms of relative coverages and tested their modality at aforementioned biologically relevant values. Ancient clones (EN and EEN) possessed relatively smooth distributions of relative normalized coverages with peaks close to 1, suggesting gene conversions as the main mechanism causing their LOH events. In contrast, recent clones (ET, EET and ETT) had additional peaks located at lower values (Fig. 4a-e), indicating simultaneous operation of both processes (Tucker et al. 2013).

**Fig. 4:**
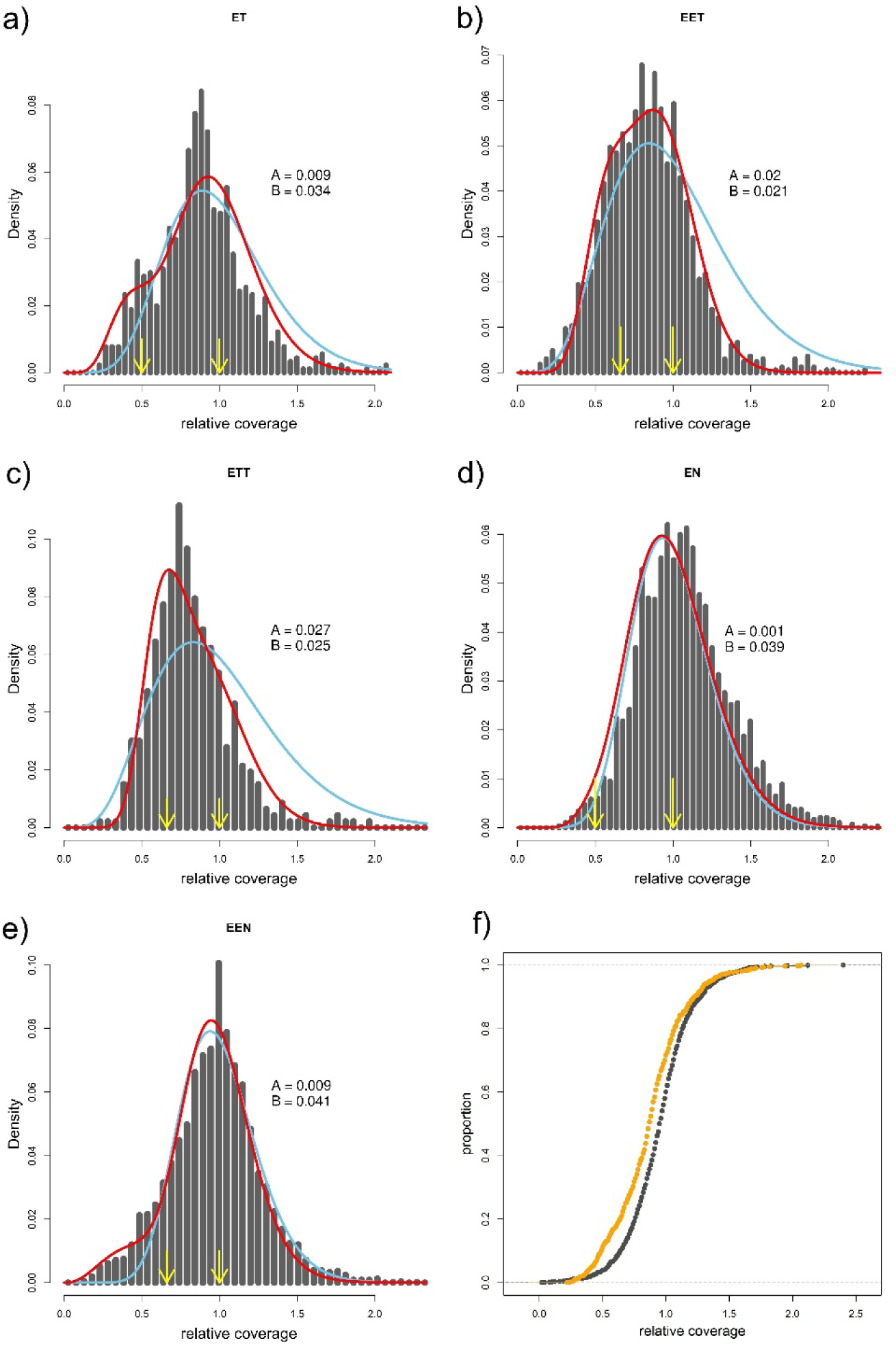
**a-e)** Histograms of relative coverages at LOH loci in ET, EET, ETT, EN and EEN biotypes, respectively. Arrows depict biologically meaningful values (for given ploidy); blue lines represent the fit of single Gamma distribution with mean centered at value 1, red lines represent the fitted mixture of two Gamma distribution with the coefficients A and B indicating proportions of both Gamma distributions in the combined model; A relates to the distribution assuming the mean relative coverage ∼1, B to the distribution with the mean ∼0.5 or 0.66 (for diploid or triploid biotype, respectively); **f)** orange represents empirical cumulative distribution function (ECDF) of relative coverages at LOH sites of ET biotype, black represents ECDF of relative coverages at the same sites, but taken from parental species, where no deletions are expected. ECDF curves for other biotypes are provided in Fig. S1.

To formally test whether observed data may be explained by single process or several simultaneously operating processes, we compared the fits of each histogram by single gamma distribution as a proxy for operation of only one process, or mixtures of more distributions with means fixed at aforementioned biologically relevant values. Nonlinear least square method was used to estimate the parameters controlling relative proportions of distributions in combined models (A and B parameters control the rate of conversions / hemizygous deletions in two-gamma distribution model while A, B & C control the rates of conversions & hemizygous & double deletions in three-distribution model). The most complex model assuming the occurrence of conversions and both single and double deletions (three gamma distributions) did not significantly improve the fit to triploids’ data. However, mix of two gamma distributions assuming gene conversions and hemizygous deletions significantly outperformed any single-distribution in all datasets (F test in diploids: EN: df = {40,38}, F=2.583; ET: df = {41,39}, F=7.87; triploids: EEN: df = {44,42}, F=11.01; EET: df = {44,42}, F=112.6; ETT: df = {31,29}, F=57.49), suggesting that joint operation of both processes.

To further validate if deletions indeed occur in hybrids, we used Kolmogorov-Smirnov test to compare distributions of relative coverages at hybrids’ LOH sites with the distribution of coverages at the same sites in parental species, where no deletions are expected. This comparison also indicated that all hybrid biotypes have significant excess of low-coverage LOH sites, again suggesting that deletions exist in hybrids (see Fig. 4f and Fig. S1 for details).

Both tests thus documented the existence of LOH sites with decreased DNA content suggesting that *LOH events are caused by simultaneous operation of conversions and hemizygous deletions in all biotypes. However, double deletions in triploids are very rare or absent*.

### 3.4. Accumulation of LOH is biased with respect to parental subgenome, ploidy and hybrid type

We noted that LOHs were non-randomly distributed among hybrids, and there were several trends behind such unevenness. First, there was a clear bias in retention of parental subgenomes (Fig. 3b). In triploids, virtually all LOH sites detected possessed allele of that parent which contributed two chromosomal sets. This type of bias in triploids is likely methodological, since losses of one allele of the diploid subgenome would still appear heterozygous and hence escape our attention. However, significant bias was observed in diploid hybrids with ***preferential retention of C. elongatoides allele*** at ∼80% LOH sites in ET and ∼87% in EN hybrids.

Second, the ***hemizygous deletions were significantly more common in triploid hybrids than in their diploid counterparts***. Specifically, the A / B ratio of combined Gama distributions suggests that deletions accounted for only ∼21% LOH events in ET diploid hybrids, while their contribution rose to ∼ 50% in triploid EET and ETT hybrid forms ; Fig. 4a-c. Similarly, in *elongatoides-tanaitica* hybrids, deletions accounted for less than 0.1 % of LOH events in diploid EN hybrids, while triploid EEN possessed ∼18 % of deletions at LOH sites ; Fig. 4d-e. These differences appeared highly significant after comparing data fitting by ‘free’ mixed gamma model using optimized A / B ratios and by ‘forced’ model with A / B ratio fixed to values estimated from triploids (EN with A / B estimated from EEN data: df = {40,38}, F=2.33; ET with A / B estimated from EET data: df = {41,39}, F=14.14).

Finally, ***hemizygous deletions were more common among recent asexuals than in ancient clones***, as evident from comparisons of A / B ratios between recent (ET) and ancient (EN) diploid clones (∼21% vs. ∼0.1%) as well as of recent (EET or ETT) and ancient (EEN) triploid clones (∼50% vs ∼18%); Fig. 4a-e.

### 3.5. Occurrence of LOH is related to sequence composition, allelic expression and gene function

#### 3.5.1. LOH depends on GC content but patterns are complex

To investigate potential GC bias, we first inspected transcriptome-wide GC contents of sexual and asexual forms, measured either across all positions or only at the relatively neutral third codon positions, but found no significant differences between any biotypes. Next, we performed a more fine-scale analysis on E-TN diagnostic positions and separated all detected LOH sites into E-like or TN-like groups, depending on parental allele retained. Comparing parental sequences with each hybrid we inferred how many LOH sites underwent A/T → G/C substitutions, G/C → A/T substitutions or no change in GC content (i.e. A ↔ T or G ↔ C substitutions). We used contingency tables to compare these counts with overall A/T - G/C differences between respective parental species across all E-TN diagnostic positions and found that E-like LOH events were significantly biased in favor of A/T → G/C substitutions in triploid (EET, EEN) and diploid (ET, EN) biotypes. This bias was ∼25% on average and proved significant in every individual after FDR correction. In contrast, we observed no such GC bias in TN-like LOH sites of any biotype (Fig. 5a). Our data thus indicate that ***GC-dependence has complex background and occurs only during loss of taenia/tanaitica allele, but not in the opposite direction***.

**Figure 5:**
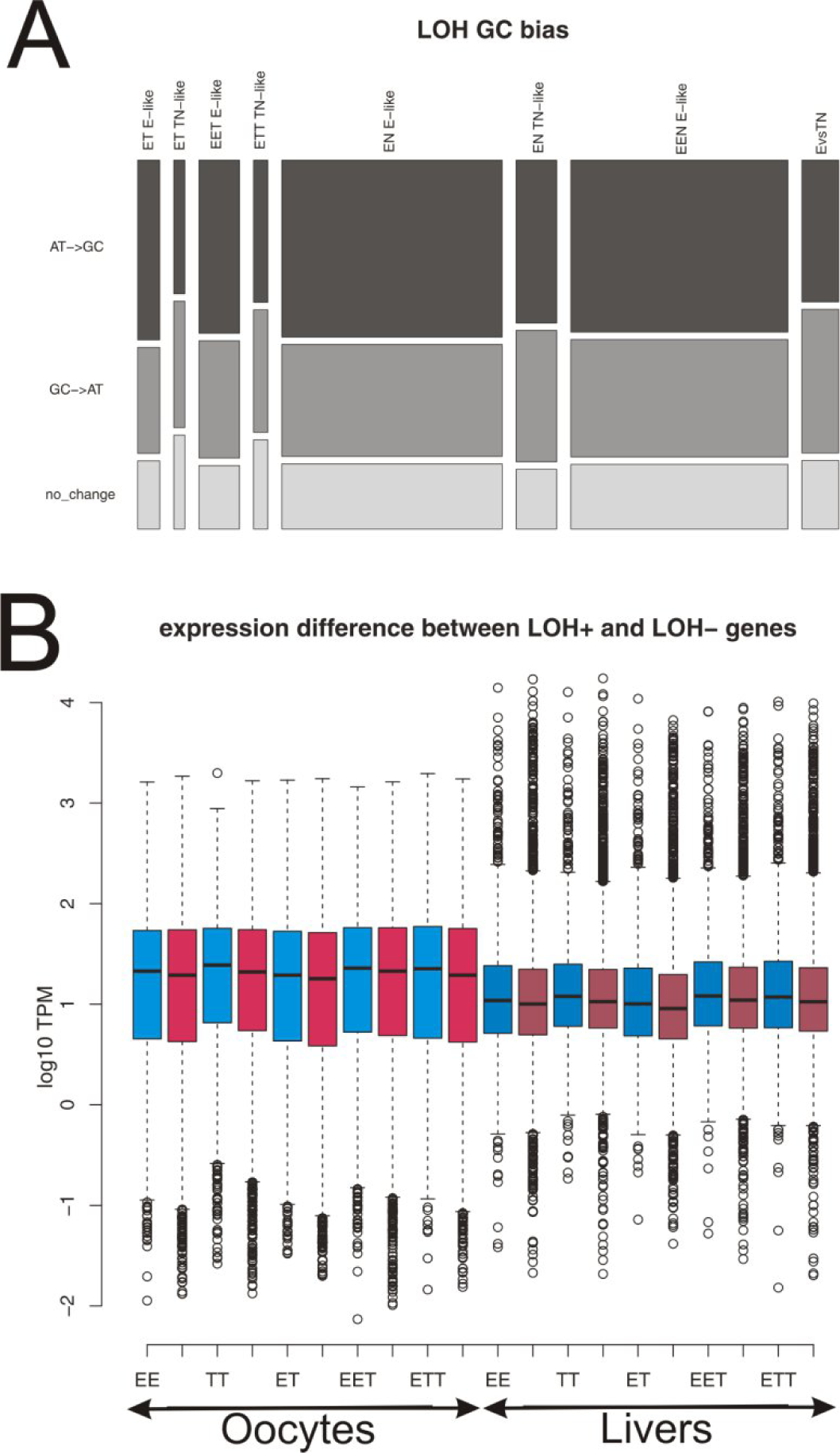
A: G/C bias at asexuals’ LOH sites as demonstrated for one representative of every biotype. Each individual is represented by two columns with E-like (left) and T-like (right) LOH events. Bar widths scale with absolute numbers of observed LOH events in each individual, while heights of color fields demonstrate proportions of LOH causing weak to strong, strong to weak and no GC change. The last bar (E vs TN) represents overall differences between C. *elongatoides* and both other parental species at all E-TN diagnostic sites. Note that in E-like LOH events, all hybrids show consistent and significant increase in weak to strong substitution rates as compared against interparental divergence. No such shift has been detected at TN-like LOH sites. B: TPM normalized expression characteristics of LOH positive (blue) and LOH negative (red) genes in all *Cobitis* biotypes analyzed by Bartoš et al. 2019. For better orientation, the oocyte data (left part) are depicted in lighter color tones, while liver data (right part) are in darker tones.

#### 3.5.2. LOH is affected by gene’s transcription

We evaluated the effects of allele expression on LOH occurrence using recently published transcription profiles of livers and oocytes in sexual (*C. elongatoides*, *C. taenia*) and asexual (ET, EET, and ETT) females (Bartoš et al. 2019). While Bartoš et al. analyzed different individuals, they used the same reference transcriptome and thus we could investigate the transcription profiles of those genes which carried E-T diagnostic SNPs and were either LOH-positive (E-like or T-like LOH bearing), or LOH-negative, according to present exome-capture results. We applied two approaches to test for differences in gene and allele expression between these categories.

First, we found that ***LOH events tend to occur in genes with abnormally high expression levels.*** Specifically, we compared the TPM-normalized (transcript per million) expression levels of Bartoš et al.’s (2019) data and found that the genes where present analysis discovered LOH events had significantly higher expression levels in liver tissue of all biotypes (FDR corrected WMW test p values < 0.02 for all biotypes). Same trends, albeit insignificant, were observed in oocytes.

Second, we compared published DeSeq2-normalized expression levels of both parental species to test if less expressed alleles tend to be preferentially lost during LOH events. We adopted such an indirect approach of comparing both parental species instead of directly inspecting allelic expressions in hybrids because Bartoš et al.’s (2019) data in hybrids might have been affected by undetected allelic deletions. Moreover, the interparental expression divergence is a good predictor of relative allelic expression given pervasive cis-regulation between subgenomes (Bartoš et al. 2019).

We found that distributions of *C. elongatoides* / *C. taenia* log_2_ fold change (log2FC) at genes bearing E-like LOH were very similar to those of LOH-negative genes (oocytes: mean log2FC -0.22 vs -0.21; livers: mean log2FC -0.14 vs -0.16; note that prevalence of negative values is in agreement with slight but pervasive under-expression of *C. elongatoides* documented by (Bartoš et al. 2019). In contrast, log2FC values were more negative (i.e. deviated towards *C. taenia*) in genes bearing T-like LOH than in LOH-negative genes (oocytes: mean log2FC -0.34 vs -0.22; livers: mean log2FC -0.22 vs -0.14). Kolmogorov-Smirnov test proved that such a difference was significant in oocytes (p value = 0.022), suggesting that the ***fate of hybrid’s alleles is affected by expression levels*** inherited from parental species. However, ***the effect is again not symmetrical with respect to subgenome ancestry,*** since less expressed alleles are preferentially removed during E-like LOH events while no bias was observed in the reciprocal direction.

#### 3.5.3. LOH accumulate in genes with specific functions

Finally, we investigated whether LOH events accumulate in genes with specific functions. For this purpose we performed two tests on loci with diagnostic SNPs that were successfully annotated. First, as a proxy for selection regime of particular genes, we used *d*_N_/*d*_S_ values among orthologous sequences of *C. elongatoides*, *C. taenia* and *C. tanaitica* published by (Kočí et al. 2020) and found in both *elongatoides*-*taenia* and *elongatoides*-*tanaitica* hybrid types that ***LOH-positive genes were characterized by significantly lower dN/dS values than LOH-negative genes*** (Linear mixed effect model with pairs of individuals as random factor; LRT p value for *elongatoides*-*taenia* = 1.604095e-135; p value for *elongatoides*-*tanaitica* = 1.518951e-05).

Next, we searched if LOH positive genes are associated with particular Gene Ontology terms using the GO::TermFinder (Boyle et al. 2004). Results are shown in Table 1 and Table S3, which list top 20 enriched GO terms of each category in each biotype. It shows that GO terms associated with membrane coats and endoplasmatic reticulum were enriched among LOH-positive genes in EET, EN and EEN biotypes with p-values corrected for multiple tests below alpha level 0.1. It also shows that some GO terms, whose corrected p value exceeded the threshold level, were repeatedly encountered among top enriched GO terms in several hybrid biotypes including independently arisen *elongatoides*-*taenia* and *elongatoides*-*tanaitica* hybrid types. These namely contained Cellular compartment type GO terms associated with cell-cell junction and cell-substrate adherence junction and Biologic processes type GO terms of cellular biogenic amine metabolic process and Protein glycosylation.

**Table 1:**
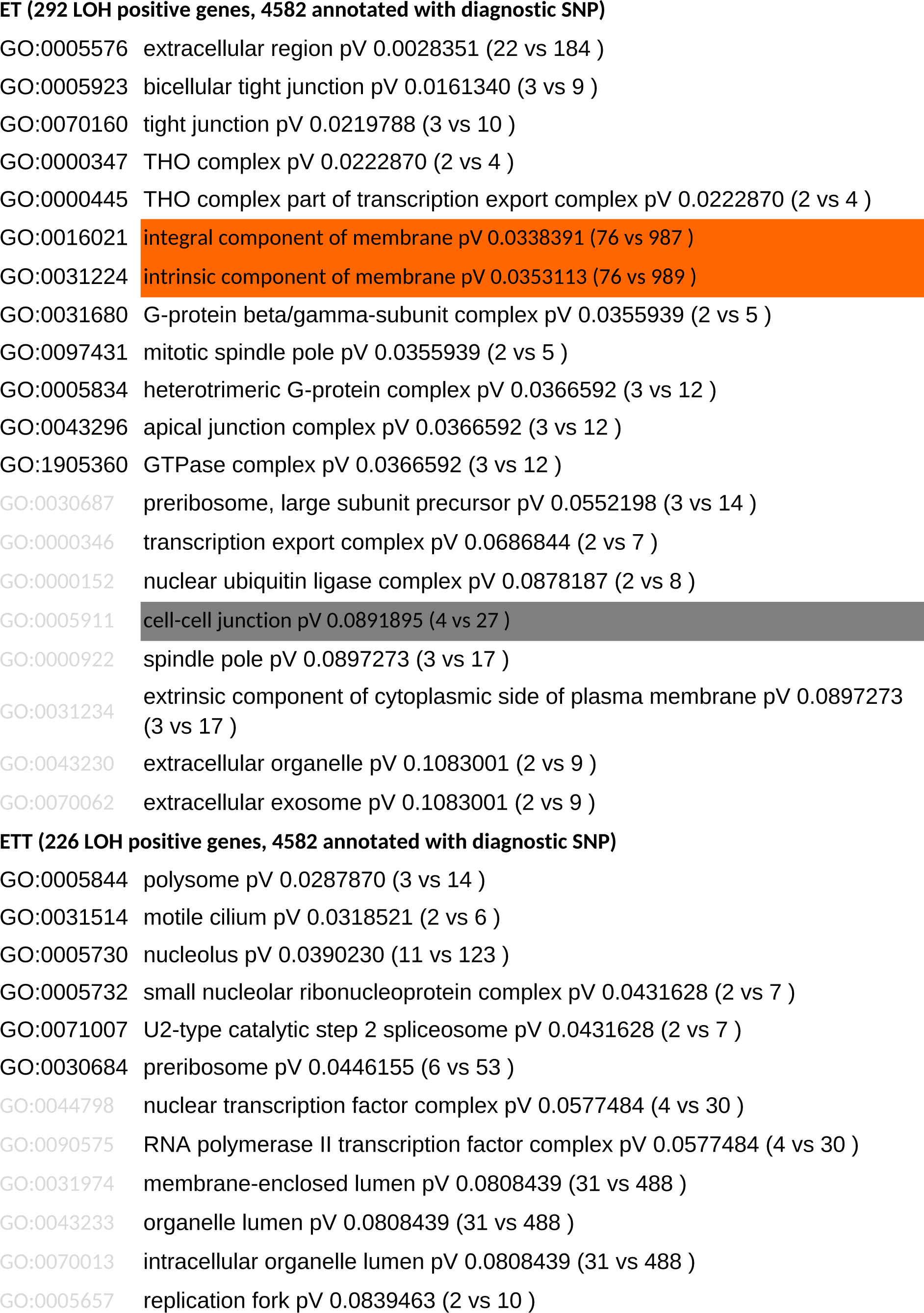
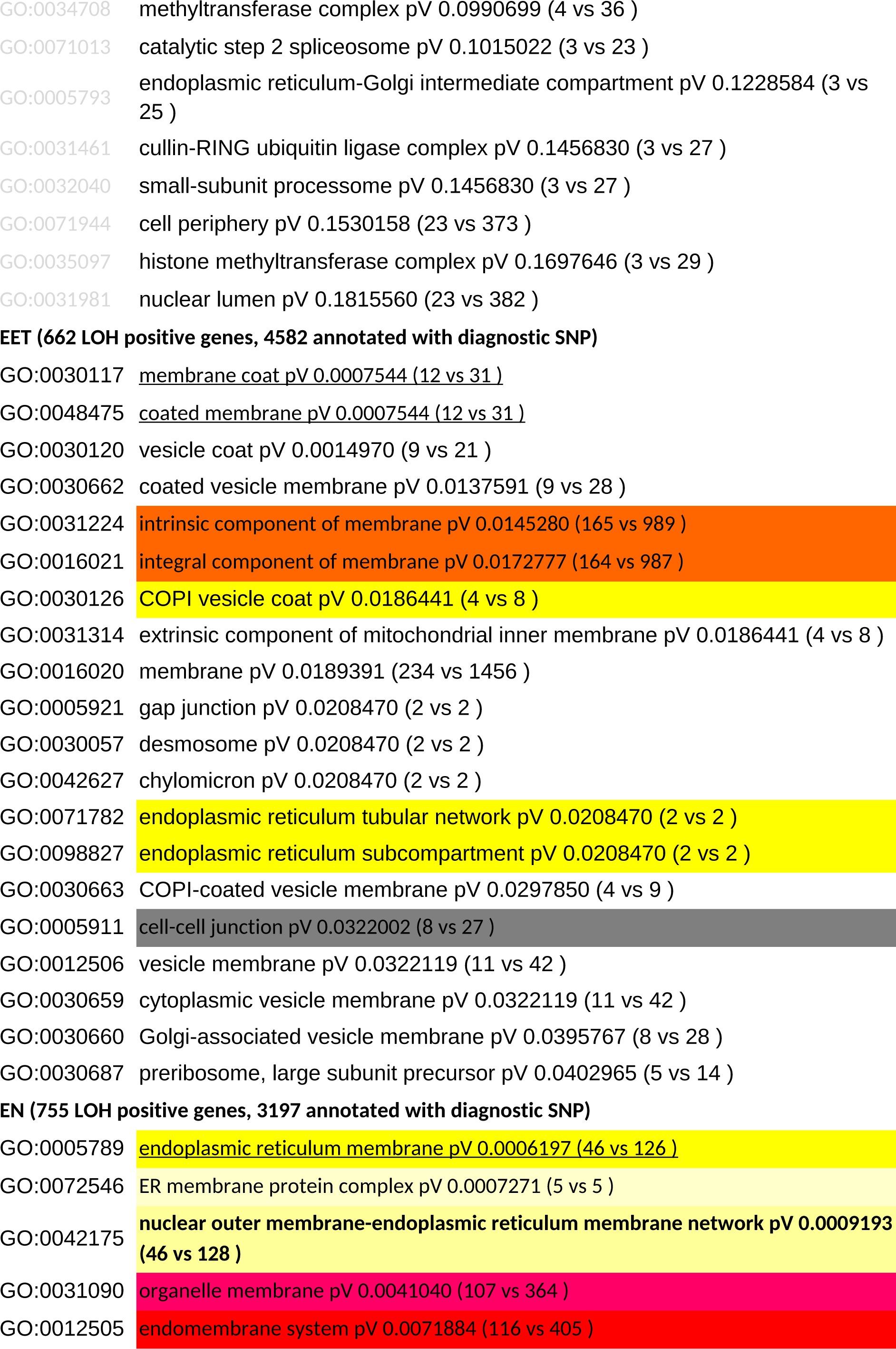
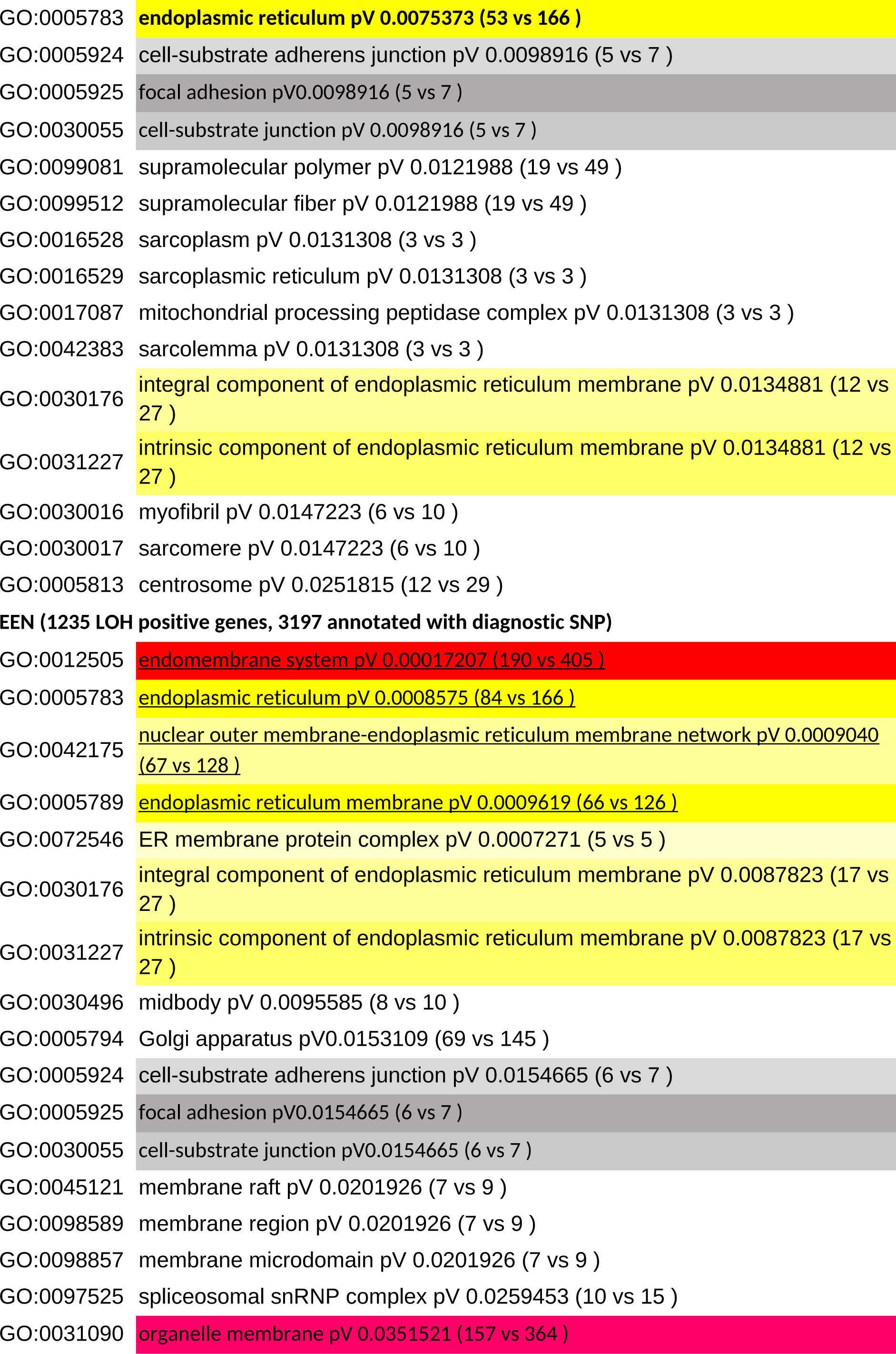
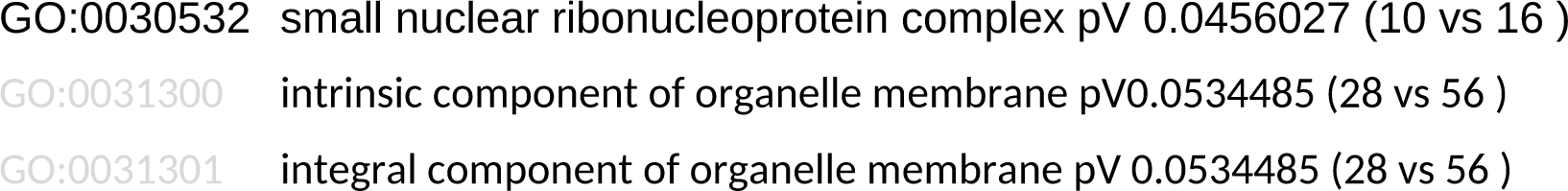
Top 20 enriched GO terms in cellular compartment GO category ranked by p value. For each hybrid biotype we indicate number of genes affected by LOH event and total number of annotated genes with diagnostic SNP relevant for given combination of parental species. For each GO term, we indicate its ID, description and uncorrected p value, as well as numbers of LOH-positive genes and total number of genes with given GO in parentheses. Underlined GO terms are significant after correction for multiple test at alpha level=0.1. Colors are used to highlight GO terms shared between distinct biotypes so that the same color across biotypes indicates GO terms that are identical or nested.

## 4. Discussion

Genomes of asexual organisms may evolve in various ways, ranging from fast restructuration to long-term conservation of heterozygosity. Such diversity of patterns may reflect the variety of gametogenetic pathways used by such organisms and supposedly, hybrids employing premeiotic endoreplication, such as *Cobitis*, can maintain integrity of the parental subgenomes without any chromosomal-scale restructurations (Majtánová et al. 2016). However, the present study showed that despite apparent stasis on a large scale, the heterozygosity gained during original hybridization may gradually deteriorate by small-scale restructurations that affect genomic regions in relation to allelic origin, sequence composition, and gene expression.

### 4.1. Large-scale stasis vs. small-scale dynamics of asexual genomes

After initial merging and duplication, allopolyploid organisms appear to deduplicate their genomes prominently via fractionation and deletions of orthologs (Yoo et al. 2014; Cheng et al. 2018; Du et al. 2020). However, we found that deletions accounted for a rather minor fraction of restructuration events in polyploid loaches and especially in diploid hybrids. A majority of LOH sites had relative coverage close to 1, thereby indicating higher incidences of recombination between orthologs. Recombination may be followed by crossover (CO), which is expected to cause long stretches of LOH spanning till another recombination site or until the telomeric ends of the paired chromosomes (Fig. 1a). However, as the cytogenetic study by (Majtánová et al. 2016) ruled out any large-scale exchange of chromosomal arms between subgenomes, it appears that most LOH events detected in this study were caused by gene conversions without COs.

Interestingly, it is unclear how such conversions between orthologs may arise since organisms employing premeiotic endoreplication should rather form bivalents between sister copies of the duplicated homologs (Lutes et al. 2010; Arai and Fujimoto 2013; Dedukh et al. 2020) (Fig. 1a). In theory, they may result from errors in homology search during early meiosis if ExT bivalents are formed, but this explanation is unlikely for two reasons. First, *C. elongatoides* and *C. taenia* karyotypes are so divergent that most orthologous chromosomes may not form proper bivalents, leading to sterility of hybrid forms that lack endoreplication, typically males (Dedukh et al. 2020). Hence, even if ectopic ExT pairings occur in cells of asexual females, the formation of proper bivalents would be unlikely and gametes would not be formed. Second, (Dedukh et al. 2020) documented the occurrence of true COs in ExE and TxT bivalents in hybrid females (Fig. 1a) suggesting that hypothetical ExT bivalents would result in the exchange of large pieces of chromosomal arms, which was not observed (Majtánová et al. 2016). An alternative explanation would, therefore, assume the role of mitotic conversions, which are important in DNA damage repair (Helleday 2003). Indeed, mitotic conversions have been hypothesized to impact the evolution of asexuals (Omilian et al. 2006; Mandegar and Otto 2007), although, to our knowledge, they have not yet been directly observed in any multicellular asexual organism.

Whatever the underlying mechanism, the fact that LOH sites are heritable and shared among clone-mates suggests that LOH events occur in the germline. The genes affected by LOH events also clearly possess some characteristics typical of loci undergoing conversions. Namely, LOH-positive genes have above-average expression levels, which is consistent with the hypothesis that DNA of transcriptionally active loci is more relaxed and, hence, prone to double strand breaks (DSB), followed by repair cascade, including the recombination machinery (González-Barrera et al. 2002; Cummings et al. 2007). *Cobitis* hybrids also tend to replace the less expressed parental allele by the more expressed allele, which is in-line with growing evidence that more transcribed homoeologs are preferably utilized as templates during DSB-induced gene conversion (Schildkraut et al. 2006). Finally, the prevalence of AT → GC substitutions on some LOH sites conforms to the expected GC bias in template preference (Duret and Galtier 2009; Williams et al. 2015).

Interestingly though, our study revealed that ***processes affecting asexual genomes are strongly dependent on the ancestry of the allele acting as a template***. Namely, the preferential retention of the more transcribed allele was apparent only during *elongatoides → taenia* allele replacement (T-like LOHs), whereas GC bias was detected only in the opposite direction (E-like LOHs). However, overall GC contents were not notably affected by these processes as we found no differences between sexual and asexual forms at the transcriptome-wide scale. This suggests that the predicted increase in GC content (Bast et al. 2018) cannot be generally applied to all types of asexuals and the ancestry of subgenomes should be taken into account.

### 4.2. Impact of LOH on evolution of hybrids and polyploids

Genome rearrangements may bring both, the benefits as well as the constraints to the asexual organism, and their accumulation may be facilitated by the lack of requirement of proper homology for chromosomal pairing (Sunnucks et al. 1996). Consequently, conversions & and deletions of genes or even chromosomal arms may potentially proceed at faster rates than mutation accumulation in some asexuals (Triantaphyllou 1981; Sunnucks et al. 1996; Normark 1999; Spence and Blackman 2000; Tucker et al. 2013). Hence, they may slow mutational deterioration (e.g., Muller’s ratchet process) by erasing deleterious mutations or increasing the fixation rate of beneficial mutations (Khakhlova and Bock 2006; Mandegar and Otto 2007). However, recombination *per se* may have mutagenic effects on its own (Arbeithuber et al. 2015).

In any case, recent analysis of mutation accumulation and fitness deterioration proposed several reasons why LOH events do not play important role in slowing the Muller’s ratchet in asexual loaches (Kočí et al. 2020). In brief, approximately only <1.5% of private asexual SNPs occur in homozygous states, indicating a rather efficient mechanism of clonal reproduction, when a majority of newly acquired mutations occur in heterozygous states on one chromosome with little possibility of recombination or conversion. Consequently, the observed rate of LOH accumulation was low occurring in only approximately 10% genes after ∼300 kya of evolution in the oldest clone. This is orders of magnitude less than that in aforementioned taxa, wherein such processes have been hypothesized to interfere with the accumulation of deleterious mutations. Finally, the efficiency of LOH in mutation erasing should increase with clonal age as rare LOH events would likely happen on genes where mutations have not yet accumulated in recent clones, whereas the emerging LOH event has a higher chance to “heal” previously mutated genes in older clones. However, such a process should lead to exponential correlation between the proportions of LOH and private asexual SNPs, which was not observed by Kočí et al. (2020). This does not indicate that LOH events may not counteract the ratchet in some asexuals; however, their role in removal of deleterious mutation is supposedly smaller in organisms with relatively efficient clonal reproduction, such as *Cobitis*.

Nevertheless, LOH events considerably impact the evolution of the studied asexuals by other mechanisms, as will be discussed in following paragraphs.

#### 4.2.1. Effects of deletions are less severe in polyploids

Hemizygous deletions cause aneuploidies on sub-chromosomal levels and may, therefore, modify the stoichiometry between interacting components of molecular complexes (Birchler and Veitia 2012) or between transcription factors and their binding sites, thereby affecting gene regulation (Veitia et al. 2013). The magnitude of their effect probably scales with the length of the deleted genomic region, i.e., long deletions affecting many genes have stronger effect than short-range deletions (Veitia et al. 2013). This may explain why *Cobitis* hybrids retained stable karyotypes with no chromosomal-scale deletions (Majtánová et al.

2016), but small-scale deletions of individual genes do occur and are not removed by selection. Veitia et al. (2013) further postulated that the impact of aneuploidy should depend on the allelic dosage. Consequently, hemizygous deletions would have weaker effects in triploids (changing allelic copy numbers from 1 to 2/3, which is relative to the rest of genome) than those in diploids (changing from 1 to 1/2), and double deletions in triploids (change changing from 1 to 1/3) would have the most severe effects. This may explain why triploids possessed higher proportion of hemizygous deletions than their diploid counterparts, but in the same time, we found no evidence of double deletions.

The hypothesis that allelic deletions have a mostly negative impact may also explain why young clones possess a relatively higher proportion of deletions than old clones. Indeed, selection-based removal of deleterious mutations requires some time proportional to the selection coefficient, population size, and genetic background, and hence, although recent clones acquired lower absolute numbers of LOH events, they would have a higher fraction of deletions due to a time-lag necessary to remove these deleterious mutations (Johnson and Howard 2007). In fact, we observed similar differences between young and old clones with regard to relative loads of nonsynonymous mutations (Kočí et al. 2020).

#### 4.2.2. Biased genome fractionation and template-preference

Another prominent pattern was the strong preference for *elongatoides* subgenome retention at LOH sites (Fig. 3b). Biased genome fractionation is commonly observed among hybrids and allopolyploids and may have various explanations, ranging from mechanistic reasons, when one ortholog induces the other’s loss, to natural selection, preferring the fixation of one allelic type in hybrid populations. For instance, fractionation bias has been put in context to subgenome expression dominance in hybrids (Yoo et al. 2014; Alexander-Webber et al. 2016; Cheng et al. 2018). Mechanisms causing such expression dominance are unclear and may relate to various processes such as cis-/trans divergence, unequal content of transposable elements, or levels of heterochromatinization among parental species (Woodhouse et al. 2014; Bottani et al. 2018). In any case, it has been proposed that once expression dominance occurs, loss of homoeologs from the lower-expressed subgenome would be preferred by selection due to less severe consequences (Yoo et al. 2014).

Interestingly, our data contrast this prediction, as the preferentially retained subgenome—*elongatoides*—was clearly not dominant in hybrids. Instead, Bartoš et al.’s (2019) data showed significant bias toward *taenia*-like expression of ecologic and phenotypic traits and overall expression level dominance of the *taenia* subgenome in hybrid transcriptomes with slight total prevalence of *taenia* transcripts in somatic tissue (∼1.5%) and germline (∼4%) of diploid hybrids. This suggests that ***expression dominance is not the causal explanation for biased genome fractionation*** in *Cobitis* hybrids.

Selective elimination of one parental subgenome may be particularly adaptive in gynogens by increasing their similarity to the parental species that provides them with the sperm, thereby increasing the chance to be fertilized (Beukeboom and Vrijenhoek 1998). Interestingly, all investigated ET hybrids coexist with *C. taenia*, ***making it unlikely that the preferential loss of C. taenia alleles provides such type of sex-mimicry*.**

Our data, thus, suggest that the ***causal link between transcriptome-wide expression dominance and biased genome fractionation is more complex*** than that predicted by the aforementioned hypotheses. For instance, Bartoš et al. (2019) documented that magnitude of *C. taenia* expression dominance differs between somatic traits and germline, suggesting that ***biased genome fractionation may reflect tissue-specific expression characteristics*** and other traits that often escape a researcher’s attention.

We may also speculate that biased retention of the *C. elongatoides* subgenome may reflect its special “mechanistic” properties. Several reasons for biased template preference have already been proposed, including different expression levels of orthologous alleles (Schildkraut et al. 2006), different GC contents (Duret and Galtier 2009; Williams et al. 2015), or the effect of maternal ancestry when maternal endonuclease systems may preferentially induce DSB on paternal chromosomes, thereby causing biased DSB repair and unequal gene conversion (Wang et al. 2010). However, none of these hypotheses may sufficiently explain the prevalence of E-like LOH as the *C. elongatoides* subgenome possesses neither higher expression levels nor different GC content and the maternal ancestor of all studied *elongatoides*–*taenia* hybrids was *C. taenia*. The observed bias may thus reflect other phenomena, such as specific distribution of epigenetic markings and methylation, which are known to affect recombination landscape (Mirouze et al. 2012), and may acquire unexpected and non-additive patterns in hybrids compared to their parents (Hegarty et al. 2011).

Nevertheless, although we did not identify the proximate reason for biased subgenome retention, our data do indicate that both subgenomes differ in their ability to induce LOH events in hybrids. For instance, we observed a higher proportion of LOH events in EET triploids than those in ETT triploids (Fig. 3). Given that detection of LOH events is limited in triploids by the presence of two conspecific allelic copies, a higher fraction of allelic loss/replacements would escape our attention in ETT than in EET triploids if E-like LOH occur more frequently than T-like LOH. Moreover, we already mentioned that E-like and T(N)-like LOH events differ with respect to allelic expression or GC bias, ***suggesting some fundamental differences between hybrid subgenomes in the ability to induce allelic loss or replacement***.

### 4.3. LOH preferentially accumulates in particular gene pathways

Benefits of LOH in hybrids has been directly tested in only a few studies (Smukowski Heil et al. 2017; Smukowski Heil et al. 2019). Even in these cases, the benefits of LOH appeared quite complex and specific for a given allele, subgenome, and environmental conditions (Lancaster et al. 2019). Although we could not directly examine the effects of LOH, our study ***revealed some parallel trends in LOH among independent hybrid strains.*** This suggests that LOH tends to accumulate in genes characterized by some common expression and functional properties, such as higher-than-average expression levels and lower-than-average *d*_N_/*d*_S_ ratios, indicating strong purifying selection to maintain their functionality.

Furthermore, we found that some gene pathways appear to be more affected by accumulation of LOH events than others, as apparent from the significant enrichment of GO terms associated with endomembrane systems, endoplasmic reticulum, and coated vesicles in some biotypes (Table 1). We have no explanation for such observation at the moment, but it is worth mentioning that genes with these GO terms often participate in multimeric protein complexes, ensuring vesicle tethering, coating, and transport to membranes. As subunits of such complexes are coevolving to maintain proper functionality, it is tempting to speculate that hybrids would profit from removal of heterozygosity because a mix of protein interactors from diverged orthologs may negatively impact composition of the entire complex.

Unfortunately, the power of GO analysis was weakened by the relatively low number of annotated genes with diagnostic SNPs in a manner that some GOs appeared insignificant after p-value correction, albeit all their genes carried LOH in some biotypes (Table 1).

Interestingly, same or nested GO terms were sometimes recorded among top-enriched GO terms of different hybrid biotypes, thereby hypothetically indicating LOH accumulation also in other gene pathways, which would, therefore, be worth studying further, e.g., the concerned GO terms associated with cell–cell junction; cell–substrate adherence junction, containing genes expressed since the early zygotic embryogenesis that take part in cell migration; and cell–cell communication (Siddiqui et al. 2010; Goonesinghe et al. 2012; Matsui et al. 2015).

## Conclusions

There is increasing evidence suggesting that genomes of asexual, hybrid, and polyploid taxa evolve dynamically with selective filtering of parts of the parental subgenomes. Some common trends have been identified in genome evolution of unrelated hybrid/polyploid taxa, e.g., in relation to gene/allelic expression. Although patterns revealed in the present study were generally consistent with several previously reported trends, our data do not support some widely cited ideas, such as preferential retention of transcriptionally dominant subgenome. The study on asexual hybrid loaches, thus, revealed that genome fractionation is a very complex process involving simultaneously operating mechanisms that range from a priori bias in template selection to selective fixation of adaptive LOH variants. Our study demonstrated that the relative impact of involved mechanisms likely depends on the reproductive mode, origin of particular subgenome, allelic sequence composition and transcription activity as well as on properties of involved genes and environmental conditions. In combination with recent advances in understanding the effect of aneuploidies (Birchler and Veitia 2012; Veitia et al. 2013), the data acquired on taxa, such as asexual hybrid loaches, can provide invaluable insight into the role of gene dosage in genome evolution in hybrids and neopolyploids. Investigation of genome evolution in hybrid and polyploid taxa may also provide important information about fundamental biological processes, such as meiosis and mitosis.

## 2. Material & Methods

### 2.1. Studied specimens

The study is based on exome-capture data from 46 specimens including *C. paludica* as outgroup, thee parental species *Cobitis elongatoides*, *C. tanaitica, C. taenia* and their asexual hybrids of various genomic compositions (see Fig. 1 and Table S1 for details). The specimens were a priori categorized into taxonomical units using flow cytometry and published PCR-RFLP markers (Janko et al. 2007). As in (Janko et al. 2018), we also included two laboratory *elongatoides-taenia* F1 hybrids with their parental individuals as a control of quality of base calling and LOH site detection.

### 2.2. DNA sequencing, SNP calling and identification of clonal lineages

Isolated gDNA was sheared with Bioruptor™ to proper fragment distribution, tagged by indices, pooled, hybridized to custom-designed exome-capture probes (Janko et al. 2018) and captured fragments were sequenced on Illumina NextSeq in 75 bp paired-end (PE) mode. To verify the robustness of exome-capture data and subsequent interpretation, we also performed whole-genome shotgun sequencing of single ET hybrid using the HiSeq X Ten sequencing platform in pair-end mode (average fragment length 200 bp, library preparation and sequencing performed by Macrogen®). Obtained reads were quality-trimmed by fqtrim tool (Pertea 2015); minimum read length 20 bp; 3’ end trimming if quality drops below 15 and aligned to *C. taenia* reference transcriptome that was published and cleaned from potentially paralogous contigs by (Janko et al. 2018). To identify mitochondrial variants, we also mapped the reads to published *C. elongatoides* mitochondrion (accession no. NC_023947.1). Mapping was performed with BWA MEM algorithm (Li and Durbin 2009) and resulting files were processed with Picard tools (Broad Institute 2015). Individuals’ variants were called with GATK v3.4 HaplotypeCaller tool and all individuals were jointly genotyped using the GenotypeGVCFs tool (McKenna et al. 2010; DePristo et al. 2011; Van der Auwera et al. 2013). Variant recalibration was based on available database of species-diagnostic positions (Janko et al. 2018) representing learning set for variant quality score recalibration tool VariantRecalibrator. Recalibrated variants were then filtered with ApplyRecalibration tool using 90 % tranche to filter all variants. All resulting highly confident SNPs with coverage >= 10 and genotype quality >= 20 were transferred into the relational database using our own Python3/SQL scripts.

SNP data of each specimen were subjected to clustering analysis by Plink v1.90b4 (Chang et al. 2015). To simplify the analysis, we focused solely on bi-allelic SNPs with at most two variants throughout the entire dataset. This resulted in removal of ∼1‰ of positions. We also removed 103 positions where the two laboratory F1 hybrids differed from their parents, because such variants were suspiciously present in most of the other specimens and suggested potential sequencing or demultiplexing errors rather than real variants.

To identify which hybrid individuals putatively belong to the same clone, we followed (Arnaud-Haond et al. 2007). Specifically, we created a pairwise matrix of distances between all hybrid individuals calculated from SNP mismatches. We then investigated a histogram of such pairwise distances and found a saddle point, which putatively defines a threshold distance between pairs of individuals belonging to the same or different clones.

### 2.3. Selection of species-specific SNPs, Detection of Loss of Heterozygosity (LOH) events and evaluation of their topological clustering

All SNPs that passed through aforementioned filters were attributed into one of the 10 categories according to their distribution among biotypes, following (Ament-Velásquez et al. 2016). The most important category of SNPs for this study are the species-specific variants (categories shared sh3a-b), where parental species are fixed (monomorphic) for different alleles, thereby allowing for detection of so-called Loss of Heterozygosity (LOH) events, where hybrids appear homozygous contrary to the expectation. Having detected the LOH variants in hybrids, we test whether observed LOH events tend to be randomly distributed across individual’s genes or rather tend to cluster, in which case the SNPs with LOH in given gene occur in tight proximity to each other with no discontinuation by heterozygous diagnostic sites. To do so, we identified within each gene the uninterrupted stretches of diagnostic sites with LOH and assigned each such cluster with a score (***S***) so that if the length of the LOH cluster = ***n***, then ***S=n^2***. Overall clustering score per animal is then simply represented by ***∑S***. Finally, to test whether observed clustering is nonrandom, we permuted for each hybrid individual its LOH sites across all diagnostic sites and genes, calculated ***S*** and compared simulated values to empirical scores.

### 2.4. Analysis of sequencing coverage

We calculated the normalized coverage at each LOH site of every individual using the “total read count” approach (Dillies et al. 2013) and estimated the so-called ‘relative coverage’ by dividing hybrid’s normalized coverage at given site by normalized coverages of the same site in parental species. Since deletions are unlikely in sexual species, this allows detection of hemizygous deletions in hybrids. The expected values then depend on the ploidy of given hybrid: ∼1 would indicate the same number of allelic copies indicating a conversion event, while hemizygous deletions would generate relative values ∼0.5 and ∼0.66 in diploid and triploid hybrids, respectively, while double allelic deletion in triploids would generate values ∼0.33.

To reveal whether observed LOH events in hybrids are generated by gene conversion or hemizygous deletions, or their combination, we performed two tests. First, constructed histograms of relative coverages of all detected LOH sites for each hybrid biotype and tested their modality at values biologically relevant for gene conversion (∼1) or deletion (∼0.5 in diploids, or 0.66 and 0.33 in triploids) (Tucker et al. 2013). To test whether observed distributions deviate from unimodality and to evaluate the relative contribution of gene conversion and deletion processes, we applied the nonlinear least square method implemented in Gnuplot software to consecutively fit each histogram by gamma distributions with the shape parameter *k* optimized by the fitting algorithm and mean (*μ = α/β*) fixed at aforementioned relevant values. In case of diploids, we fitted two distributions (or their mix), centered at 1 and 0.5, while in triploids we fitted three distributions (or their mixes) centered at 1, 0.66, and 0.33. Before fittings, we followed the Freedman-Diaconis rule to select the width of the bins in each histogram (Freedman and Diaconis 1981) in order to take into account the properties of particular datasets of each biotype. In case of mix of distributions, we let the fitting algorithm estimate the optimal values of A, B and C parameters describing relative contributions of individual distributions.

We then simulated how the distribution of normalized coverages should look like, were it generated by conversions only. To do so, we generated simulated dataset for each hybrid individual, where coverage values at its LOH sites were sampled from exactly the same number of sites/genes in parental individuals. As such, we obtained realistic null expectation of relative coverages while taking into account used methodology of DNA sequencing and bioinformatic treatment. Such a null distribution simulated for each hybrid biotype has been compared with the empirical one by Kolmogorov-Smirnov test. The simulations have been repeated several times for each biotype, randomizing both the loci considered as well as the specimens used per each locus.

### 2.5. Testing the effect of gene/allele expressions on LOH occurrence

To evaluate potential effects of gene/allelic expression on the occurrence of LOH, we compared the present data with those published by (Bartoš et al. 2019).

Using present gDNA data we categorized loci based on presence or absence of LOH (LOH positive or negative) and its direction (E-like or T-like). We then analyzed the RNA expression of corresponding loci in data of Bartoš et al., (2019) using two tests:

⍰ **Differences in overall gene expression:** To compare the expression levels of LOH-positive and LOH-negative genes in each hybrid biotype, we normalized original read counts by TPM method (Transcripts Per Kilobase Million) instead of DeSeq2 method used by Bartoš et al. (2019), since the TPM allowed for comparisons of multiple loci within each individual. The TPM normalization was performed according to (Mortazavi et al. 2008).
⍰ **Allelic expression:** Using expression data from Bartoš et al. (2019) we also investigated expression divergence between parental species as a proxy for relative allelic expression in each locus. Specifically, we extracted the DeSeq2 normalized estimates of *C. elongatoides* and *C. taenia* expression levels and tested whether the direction of LOH event (either E-like or T-like) is related to log2 fold change (FC) between parental species. The test was performed by comparing the distributions of log2FC values in LOH-negative genes with those of either E-like LOH or T-like LOH-positive genes using Kolmogorov-Smirnov test.

### 2.6. Analysis of Gene Ontology Term enrichment

The reference transcriptome was annotated with BLAST2GO tool v1.4.4 using GO database as of July 2019). From 20600 sequences, a subset of 13557 received BLASTx hit (e-value < 0.0001), from which 11314 was associated with significant GO Term annotation (default BLAST2GO settings). To identify GO terms potentially associated with LOH-positive genes, we performed GO enrichment analysis restricted to those genes, which possessed diagnostic sites, thereby technically allowing detection of LOH. P-values were calculated from hypergeometric distribution implemented in GO::TermFinder (Boyle et al. 2004) using the list of LOH-positive genes as a testing dataset for each biotype. Since gene ontologies terms are a part of acyclic directed graphs (parent and child terms are not independent), we also corrected obtained p values with permutation-based correction provided in GO::TermFinder.

## Acknowledgments

We would like to thank EMBL GeneCore crew for sequencing and advice on data processing. We also thank Jacek Stefaniak for preparation of map of samples and Dmytro Dedukh for photo of bivalents. The research was supported by the Czech Science Foundation grants GAČR 17-09807S and 19-21552S, the University of Ostrava grants SGS19/PŘF/2015 and SGS19/PŘF/2017, and the Ministry of Education, Youth and Sports of the Czech Republic grant EXCELLENCE CZ.02.1.01/0.0/0.0/15_003/0000460 OP RDE. Access to computing and storage facilities owned by parties and projects contributing to the National Grid Infrastructure MetaCentrum provided under the programme Projects of Large Research, Development, and Innovations Infrastructures (CESNET LM2015042), is greatly appreciated.

## Supplementary files

**Table S1:** List of samples including sample name, biotype, sex, sequencing statistics and geographical origin.

| sample | biotype | sex | no. of reads | no. of bases | mapped<br>reads | locality | site ID |
| --- | --- | --- | --- | --- | --- | --- | --- |
| csc008 | TT | m | 9328500 | 1890943172 | 7872341 | Tnya river, Ukraine | 22 |
| csc009 | EE | NA | 10674754 | 2200211351 | 9160050 | Kozačevski river, Bulgaria | 3 |
| csc011 | EET | f | 46169330 | 4546217825 | 24662190 | Odra R. sites RQ, Poland | 18 |
| csc012 | EEN | f | 36325574 | 3585572296 | 18876965 | Odra R. sites 509, Poland | 16 |
| csc013 | EEN | f | 44514804 | 4386326838 | 25987623 | Čierna voda, Slovakia | 1 |
| csc014 | EEN | f | 47084588 | 4630637949 | 27469787 | Čierna voda, Slovakia | 1 |
| csc015 | EEN | f | 50128474 | 4925558699 | 25581023 | Čierna voda, Slovakia | 1 |
| csc016 | EEN | f | 47466730 | 4675165684 | 28108072 | Čierna voda, Slovakia | 1 |
| csc017 | EEN | f | 62526316 | 6142494088 | 32394947 | Čierna voda, Slovakia | 1 |
| csc018 | EEN | f | 55335802 | 5446458549 | 30131168 | Čierna voda, Slovakia | 1 |
| csc019 | EE | NA | 44761144 | 4406019008 | 23043480 | Čierna voda, Slovakia | 1 |
| csc020 | EE | NA | 55359494 | 5451569580 | 28769818 | Čierna voda, Slovakia | 1 |
| csc021 | TT | m | 42119812 | 2985717253 | 20350974 | South Bug, Ukraine | 19 |
| csc022 | TT | f | 34584512 | 2448248483 | 16574306 | Kodyma river, Ukraine | 4 |
| csc024 | EE | NA | 38482140 | 2734317798 | 18243498 | Odra R. sites 801, Poland | 11 |
| csc025 | EE | NA | 37629664 | 2669330656 | 18231557 | Odra R. sites 801, Poland | 11 |
| csc027 | EE | m | 54542982 | 3908825666 | 28305773 | Kamcia r., Bulgaria | 6 |
| csc028 | TT | f | 32664314 | 2308852524 | 16036910 | South Bug, Ukraine | 2 |
| csc029 | EE | NA | 38364748 | 2723397165 | 18384671 | Kozačevski river, Bulgaria | 3 |
| csc033 | TT | f | 28098246 | 2010306058 | 14112070 | Krym Alma river, Ukraine | 5 |
| csc034 | TT | m | 27835994 | 1951741228 | 13518951 | Odra R. sites 705, Poland | 12 |
| csc035 | TT | m | 41334434 | 2940164538 | 20820869 | Odra R. sites 701, Poland | 10 |
| csc036 | NN | m | 31546234 | 2250940100 | 16153019 | Sucholužija, Ukraine | 8 |
| csc039 | NN | NA | 22567596 | 1603680625 | 10392416 | Kuban river, Russia | 14 |
| csc046 | EEN | f | 35381744 | 2486580209 | 16471358 | Odra R. sites RQ, Poland | 18 |
| csc052 | ET | f | 32020816 | 2249205626 | 14937219 | Odra R. sites 701, Poland | 10 |
| csc055 | EET | f | 34678947 | 2432666387 | 15941455 | Odra R. sites 510, Poland | 15 |
| csc056 | EEN | f | 32226860 | 2270486184 | 14849216 | Sluč Vachovcy, Ukraine | 20 |
| csc057 | ET | f | 31545522 | 2190478704 | 14343725 | Odra R. sites 701, Poland | 10 |
| csc067 | ET | f | 48483708 | 3434265969 | 23877555 | Odra R. sites 701, Poland | 10 |
| csc069 | ET | f | 16831044 | 1632439963 | 10815080 | laboratory hybrid | NA |
| csc070 | EE | m | 20691516 | 2009443636 | 13090582 | Olsztyn, Poland | 17 |
| csc071 | ET | f | 14455828 | 1399281976 | 8718131 | laboratory hybrid | NA |
| csc072 | TT | f | 12540166 | 1218069246 | 8555642 | Olsztyn, Poland | 21 |
| csc073 | EN | f | 24308246 | 2377911505 | 15929335 | Ghiol lake, Romania | 7 |
| csc074 | EN | f | 19470508 | 1898963852 | 12481576 | Ghiol lake, Romania | 7 |
| csc076 | EET | f | 36597710 | 3585778583 | 24666067 | Odra R. sites 510, Poland | 15 |
| csc077 | TT | NA | 18349390 | 1778582689 | 11064568 | Malaren lake, Sweden | 13 |
| csc078 | EET | f | 17368108 | 1693830781 | 10771162 | Odra R. sites 510, Poland | 15 |
| csc079 | EEN | f | 22917494 | 2231070909 | 14125361 | Odra R. sites 510, Poland | 15 |
| csc080 | TT | NA | 12337404 | 1200073223 | 7505927 | Crimea, Ukraine | 5 |
| csc082 | ETT | f | 21529004 | 2091846439 | 13242462 | Odra R. sites 704, Poland | 9 |
| csc086 | ETT | f | 20824472 | 2036604862 | 13308483 | Odra R. sites 704, Poland | 9 |
| csc087 | ETT | f | 22230674 | 2174839517 | 14061887 | Odra R. sites 701, Poland | 10 |
| csc089 | ET | f | 50135182 | 4898480713 | 27586948 | Odra R. sites 701, Poland | 10 |
| csc090 | ETT | f | 34147738 | 3331600457 | 19960304 | Odra R. sites 701, Poland | 10 |

**Table S2:** Counts of SNPs detected per each sample and category. In parentheses, we define each category by demonstrating states at SNP in asexual hybrid (H) and both parental species (P_1_, P_2_), separated by space (i.e. H P_1_, P_2_). Note that EN and EEN hybrids were classified against elongatoides—tanaitica comparisons, ET, EET and ETT hybrids against elongatoides—taenia and the two F1 hybrids against their direct parents.

| dataset | animal | biotype | MLL | pr1a<br>(xy x x) | pr1b<br>(y x x) | prh2a<br>(xy x xy \| xy xy x) | prh2b<br>(x x xy \| x xy x) | prh2b1<br>(x y xy \| x xy y) | sh3a<br>(xy x y) | sh3b11<br>(x x y) | sh3b12<br>(x y x) | sh4a<br>(xy xy xy) | sh4b<br>(x xy xy) | total SNPs | LOH proportion | diagnostic SNPs | private SNPs<br>proportion |
| --- | --- | --- | --- | --- | --- | --- | --- | --- | --- | --- | --- | --- | --- | --- | --- | --- | --- |
| elongatoides-tanaitica | csc012 | EEN | 5 | 6907 | 7 | 20054 | 23850 | 254 | 26346 | 2527 | 59 | 1195 | 645 | 81844 | 0,0894 | 28932 | 0,0845 |
|  | csc013 | EEN | 9 | 6541 | 10 | 20704 | 24035 | 202 | 26851 | 2462 | 60 | 1214 | 639 | 82718 | 0,0859 | 29373 | 0,0792 |
|  | csc014 | EEN | 7 | 7193 | 12 | 21336 | 23703 | 242 | 27108 | 2450 | 48 | 1231 | 637 | 83960 | 0,0844 | 29606 | 0,0858 |
|  | csc015 | EEN | 7 | 7084 | 10 | 21142 | 23460 | 247 | 26870 | 2460 | 48 | 1221 | 633 | 83175 | 0,0854 | 29378 | 0,0853 |
|  | csc016 | EEN | 7 | 7183 | 11 | 21291 | 23712 | 253 | 27096 | 2444 | 52 | 1229 | 636 | 83907 | 0,0843 | 29592 | 0,0857 |
|  | csc017 | EEN | 7 | 7134 | 10 | 21174 | 23652 | 252 | 27063 | 2454 | 60 | 1237 | 641 | 83677 | 0,0850 | 29577 | 0,0854 |
|  | csc018 | EEN | 10 | 5943 | 8 | 20400 | 24294 | 511 | 25820 | 3748 | 50 | 1266 | 599 | 82639 | 0,1282 | 29618 | 0,0720 |
|  | csc046 | EEN | 5 | 6816 | 7 | 19987 | 24922 | 285 | 26526 | 2682 | 67 | 1134 | 669 | 83095 | 0,0939 | 29275 | 0,0821 |
|  | csc056 | EEN | 8 | 6149 | 3 | 19369 | 24942 | 286 | 26145 | 2609 | 109 | 1108 | 657 | 81377 | 0,0942 | 28863 | 0,0756 |
|  | csc079 | EEN | 5 | 7015 | 5 | 20385 | 24488 | 255 | 26926 | 2563 | 67 | 1214 | 657 | 83575 | 0,0890 | 29556 | 0,0840 |
|  | csc073 | EN | 3 | 6423 | 46 | 16840 | 28280 | 341 | 26872 | 2676 | 377 | 1086 | 792 | 83733 | 0,1020 | 29925 | 0,0773 |
|  | csc074 | EN | 3 | 6382 | 36 | 16798 | 28192 | 338 | 26797 | 2654 | 375 | 1069 | 774 | 83415 | 0,1016 | 29826 | 0,0769 |
| elongatoides-taenia | csc011 | EET | 4 | 1673 | 1 | 28139 | 25039 | 109 | 35107 | 593 | 65 | 2237 | 798 | 93761 | 0,0184 | 35765 | 0,0179 |
|  | csc055 | EET | 4 | 1581 | 0 | 27510 | 24602 | 152 | 33638 | 756 | 52 | 1882 | 717 | 90890 | 0,0235 | 34446 | 0,0174 |
|  | csc076 | EET | 4 | 1746 | 8 | 28687 | 26460 | 100 | 36596 | 526 | 78 | 2161 | 835 | 97197 | 0,0162 | 37200 | 0,0180 |
|  | csc078 | EET | 4 | 1530 | 2 | 26096 | 24629 | 131 | 33012 | 683 | 58 | 1969 | 740 | 88850 | 0,0220 | 33753 | 0,0172 |
|  | csc052 | ET | 11 | 852 | 4 | 22318 | 30595 | 121 | 33918 | 376 | 179 | 1559 | 1008 | 90930 | 0,0161 | 34473 | 0,0094 |
|  | csc057 | ET | 1 | 1142 | 5 | 22231 | 30431 | 87 | 33495 | 371 | 175 | 1505 | 1013 | 90455 | 0,0160 | 34041 | 0,0127 |
|  | csc067 | ET | 1 | 1233 | 8 | 23668 | 31598 | 99 | 35879 | 387 | 190 | 1716 | 1078 | 95856 | 0,0158 | 36456 | 0,0129 |
|  | csc089 | ET | 1 | 1287 | 11 | 24245 | 30160 | 143 | 36624 | 454 | 177 | 2055 | 1085 | 96241 | 0,0169 | 37255 | 0,0135 |
|  | csc082 | ETT | 6 | 890 | 1 | 24687 | 28028 | 33 | 35045 | 12 | 237 | 2020 | 901 | 91854 | 0,0071 | 35294 | 0,0097 |
|  | csc086 | ETT | 6 | 912 | 2 | 24942 | 28271 | 27 | 35290 | 6 | 231 | 1970 | 912 | 92563 | 0,0067 | 35527 | 0,0099 |
|  | csc087 | ETT | 1 | 1238 | 3 | 25388 | 28038 | 30 | 35565 | 12 | 228 | 1971 | 935 | 93408 | 0,0067 | 35805 | 0,0133 |
|  | csc090 | ETT | 1 | 1278 | 3 | 26137 | 28426 | 29 | 36548 | 13 | 232 | 2153 | 947 | 95766 | 0,0067 | 36793 | 0,0134 |
| F1 family only | csc069 | ET | F1 | 40 | 0 | 6616 | 5276 | 0 | 43548 | 12 | 53 | 353 | 45 | 55943 | 0,0015 | 43613 | 0,0007 |
|  | csc071 | ET | F1 | 38 | 0 | 6378 | 5001 | 1 | 41845 | 35 | 37 | 340 | 38 | 53713 | 0,0017 | 41917 | 0,0007 |

Table S3: Top 20 enriched GO terms in Molecular Function and Biological Process GO categories ranked by p value. For each hybrid biotype we indicate number of genes affected by LOH event and total number of annotated genes with diagnostic SNP relevant for given combination of parental species. For each GO term, we indicate its ID, description and uncorrected p value, as well as numbers of LOH-positive genes and total number of genes with given GO in parentheses. Underlined GO terms are significant after correction for multiple test at alpha level=0.1. Colors are used to highlight GO terms shared between distinct biotypes so that the same color across biotypes indicates GO terms that are identical or nested.

**Figure S1:**
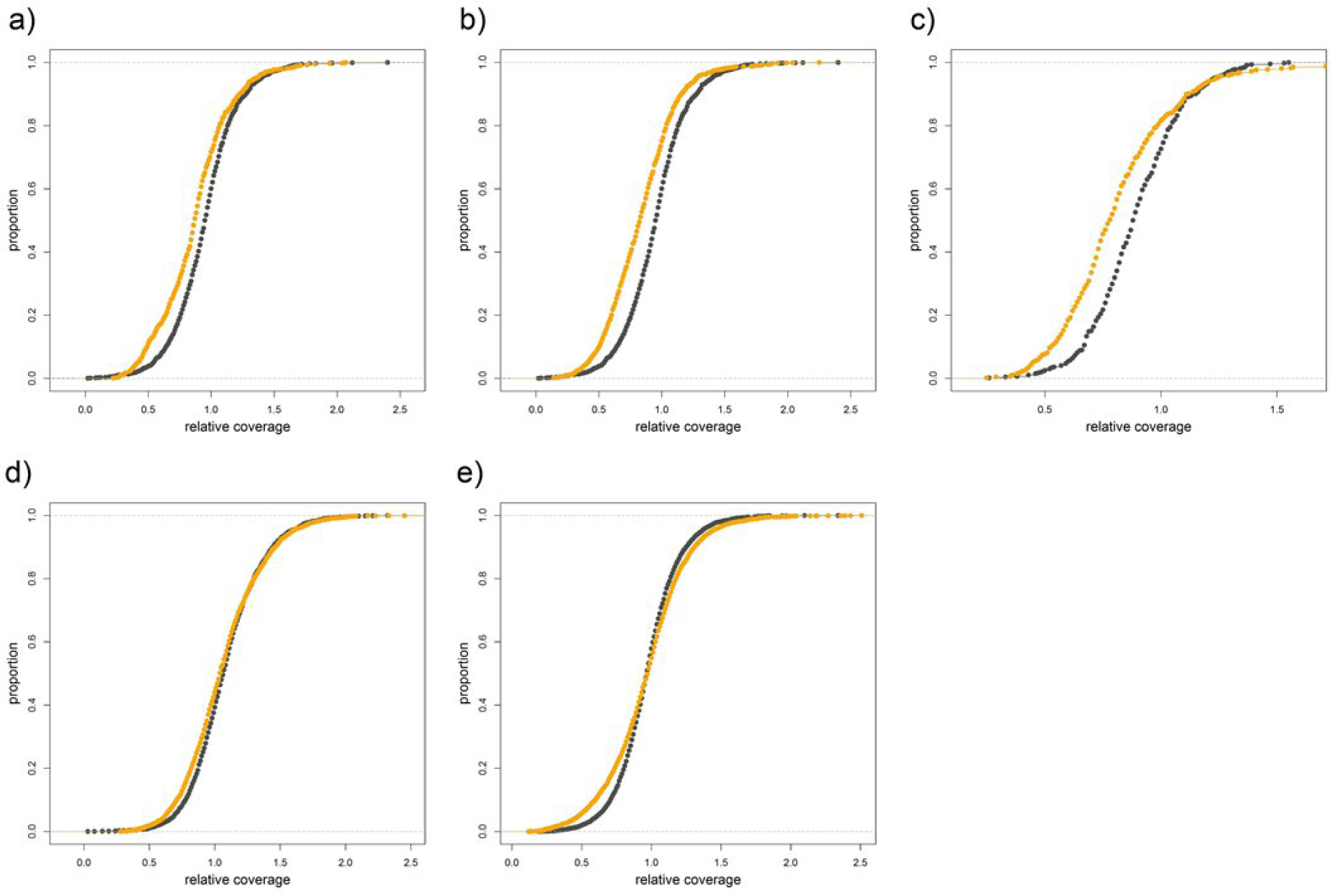
Empirical cumulative distribution functions (ECDF) for individual biotypes. Orange represents empirical cumulative distribution function (ECDF) of relative coverages at LOH sites of given hybrid biotype, black represents ECDF of relative coverages at the same sites, but taken from parental species, where no deletions are expected. Significance of differences between distributions were tested by Kolmogorov-Smirnov test. Hybrid biotypes: **a)** ET biotype, **b)** EET biotype, **c)** ETT biotype, **d)** EN biotype, **e)** EEN biotype.

